# FLOW CYTOMETRIC PROCEDURES FOR DEEP CHARACTERIZATION OF NANOPARTICLES

**DOI:** 10.1101/2024.07.18.604065

**Authors:** Valentina Tirelli, Felicia Grasso, Valeria Barreca, Deborah Polignano, Alessandra Gallinaro, Andrea Cara, Massimo Sargiacomo, Maria Luisa Fiani, Massimo Sanchez

## Abstract

In recent years, there has been a notable increasing interest surrounding the identification and quantification of nanosized particles, including extracellular vesicles (EVs) and viruses. The challenge posed by the nano-sized dimension of these particles makes precise examination a significant undertaking.

Among the different techniques for the accurate study of EVs, Flow cytometry (FCM) stands out as the ideal method. It is characterized by high sensitivity, low time consumption, non-destructive sampling, and high throughput.

In this article, we propose the optimization of FCM procedures to identify, quantify, and purify EVs and virus like particles (VLPs).

The protocol aims to reduce artifacts and errors in nano-sized particles counting, overall caused by the swarming effect. Different threshold strategies were compared to ensure result specificity. Additionally, the critical parameters to consider when using conventional FCM outside of the common experimental context of use have also been identified. Finally, fluorescent-EVs sorting protocol was also developed with highly reliable results using a conventional cell sorter.

## INTRODUCTION

The identification of extracellular vesicles (EVs) has received increasing attention in recent years. EVs are a heterogeneous population exhibiting variability in size and shape and encompassing exosomes, microvesicles and apoptotic bodies (1, 2, 3).

EVs play roles in both autocrine and paracrine signaling, functioning as mediators of cellular communication, participating in normal physiological processes as well as pathological conditions. These membrane vesicles carry proteins, nucleic acids, and bioactive lipids resembling the cell of origin (4). Once released within the extracellular space and entering the circulation, EVs may transfer their cargo to neighboring or distant cells inducing phenotypical and functional changes (5, 6). The unique characteristics of EVs identify them as non-invasive diagnostic biomarkers for the diagnosis and prognosis of various diseases, offering potential contributions to the development of next-generation therapeutic nanocarriers (7).

Among nanocarriers, virus-like particles (VLPs) also represent an attractive platform for the delivery of proteins or drugs in both vaccination and therapeutic strategies, combining ease of production, immunogenicity and high biosafety profile. Indeed, VLPs are small particles (20-200 nm) able to self-assemble, incorporating viral structural proteins and mimicking the shape, size and antigenic display of the parental virus. Importantly, since VLPs lack the genetic material, they are non-infectious and non-replicating, avoiding the risk of active infection in exposed individuals. Furthermore, VLPs can display on their surface high density of homologous or heterologous membrane-tethered glycoprotein, inducing strong cellular and humoral immune responses (8).

In this context, there is a critical need for a tool capable of distinguishing variations in the number of NPs from changes in molecular composition and providing multi-parametric analysis at the level of individual NPs.

Despite the development of various standard methods for isolating EVs from different sources, the accurate characterization and quantification of these vesicles have prompted the exploration of several technologies. Nanoparticle Tracking Analysis (NTA), Tunable resistive pulse sensing (TRPS), Dynamic Light Scattering (DLS) Electron Microscopy (EM), and Flow Cytometry (FCM) are among the technologies employed, each exhibiting its own set of advantages and disadvantages (9-13).

Although FCM is an ideal technique for high throughput, multiparametric characterization of single events, conventional FCM encounters challenges in detecting particles below the 200 nm, making precise identification of sub-micron particles difficult (14, 15).

FCM is a technique for studying single cell suspensions that is widely used in both clinical and experimental research. Its quick analysis of many events makes it a valuable tool for obtaining precise quantitative and qualitative data on samples. Employing blue laser excitation (488 nm), FCM evaluates cell morphology through forward scatter (FSC) and side scatter (SSC) signals, measuring cell size and cellular complexity, respectively. Although classically applied in immunology, hematology, oncology, and clinical biology or diagnostics, FCM is increasingly finding applications in non-conventional fields such as microbiology, industry, and the environment (16-19). Furthermore, due to its high potential, FCM applications are emerging in the identification of submicron sized EVs (20, 21).

The majority of conventional flow cytometers are designed to detect particles larger than 1 µm. However, for smaller particles, including most viruses (which range in size from 30 to 350 nm) and EVs, there is a necessity to optimise and implement analysis and acquisition protocols in order to achieve sufficient sensitivity (22, 23).

Several strategies have been employed to enhance the sensitivity of FCM. Firstly, fluorescence is essential for the detection of particles smaller than 200 nm in size (21, 23, 24). The fluorescence enables the reduction of background noise facilitating the confident identification of the EVs under investigation. Another strategy involves utilizing the 405 nm Violet Side Scatter (V-SSC) signal instead of the classic 488 nm Blue Side Scatter (B-SSC). As described by Zucker et al., V-SSC exhibits a higher separation index than B-SSC and shows a smaller coefficient of variation (CV), thereby enhancing the detection limit of FCM, allowing for the identification of 150 nm polystyrene particles (25, 26).

Although technological advances in the design of new flow cytometers have led to a significant improvement in the size sensitivity, there are still no instruments capable of clearly identifying and quantifying EVs, relying solely on physical or morphological parameters. The main difficulty is represented by the background noise, including optical and electronic noise signals, which can overlap with EVs below 200 nm in size.

Furthermore, considering the extremely small dimensions of the EVs, the swarm effect should not be underestimated. This effect occurs when two or more small particles pass through the laser at the same time and are considered as a single particle, leading to an incorrect evaluation of the number of nanoparticles present in the sample (27, 28).

The goal of our work is to develop specific and sensitive FCM procedures to identify and to quantify nanosized particles (e.g. viruses and EVs) in biological samples. To achieve this, we will use two different types of well-characterized nano-sized particles: EVs isolated from the supernatant of a melanoma cell line, and Lentivirus based Virus Like Particles (VLPs) (14, 29). Furthermore, the development of the analysis protocol for the correct identification of these particles allowed us to transfer it to cell sorter, enabling the purification of specific EVs.

## METHODS

### Cell culture

Human melanoma cells A375M and human embryonic kidney 293T cells (HEK293T) were cultured in Dulbecco’s modified Eagle’s medium. All media were supplemented with 10% heat-inactivated fetal bovine serum (FBS), 100 units/ml penicillin, 100 μg/ml streptomycin and 2 mM L-glutamine (complete media). Additional 1% MEM Non-Essential Aminoacids were added to A375M cell media. All media and supplements were from Euroclone.

HEK 293T Lenti-X human embryonic kidney cells (Clontech, Mountain View, CA, USA), used to VLPs production, were maintained in Dulbecco’s modified Eagle’s medium, DMEM (Gibco Life Technologies Italia, Monza, Italy), supplemented with 10% fetal bovine serum FBS (Corning, Mediatech Inc., Manassas, VA, USA), and 100 units/mL penicillin/streptomycin (Gibco).

Cell lines were grown at 37°C, under 5% CO2, in humidified incubators and routinely tested for mycoplasma contamination using the Mycoplasma PCR Detection Kit (ABM, #G238). All cell lines are commercially available (ATCC).

### Extracellular Vesicle labelling and collection

BODIPY-FL-C16 (C16) (4,4-Difluoro-5,7-Dimethyl-4-Bora-3a,4a-Diaza-s-Indacene-3-Hexadecanoic Acid (ThermoFisher, #D3821) or BODIPY 558/568 C12 (C12) (4,4-Difluoro-5-(2-Thienyl)-4-Bora-3a,4a-Diaza-s-Indacene-3-Dodecanoic Acid) (ThermoFisher, #D3835) were complexed with fatty acid-free bovine serum albumin (BSA, Sigma, #A8806) as described previously (Coscia et al., 2016). To isolate EVs, 60-70% confluent monolayers of A375M cells in exponential growth were incubated with 7 μM C16 or C12 at 37 °C for 4 h in medium containing antibiotics and glutamine and supplemented with 0.3% FBS (cell labelling medium). To obtain double labelled C16/C12 EVs, A375 cells were incubated with cell labelling medium containing 7 μM C16 and 7 μM C12 for 4 h. To obtain double labelled C16/Ceramide EVs, A375 cells were incubated with 5 μM BODIPY TR Ceramide (ThermoFisher, # D7540) complexed to BSA for 90 min on ice, washed twice with Hank’s balanced salt solution (HBSS) and further incubated with 5 μM C16 in cell labelling medium for 90 min at 37°C. To obtain double labelled Nef^mut^-GFP/C12 EVs, HEK293T cells were transfected with a plasmid encoding Nef^mut^-GFP as described in Ferrantelli et al. (24). Briefly, HEK293T cells were transfected with 1 µg/ml plasmid DNA complexed with 3 µg/ml branched Polyethylenimine (PEI) (Sigma-Aldrich # 408727). After 18 h, cells were washed once in HBSS and incubated with 7 μM C12 in cell labelling medium for 4 h at 37°C. At the end of the incubation with fluorescent lipids, cells were washed twice with Hank’s balanced salt solution (HBSS) containing 0.1% (w/v) fatty acid-free bovine serum albumin (H-BSA) to remove excess probe and further incubated for 24 h in complete culture medium. The conditioned medium was either immediately processed for EVs isolation or stored at 4°C for up to one week. Cells were detached from the plates with trypsin/EDTA, and their viability was assessed by Trypan Blue exclusion. Conditioned medium containing fluorescent EVs was serially centrifuged at 2000g for 20 min at 4°C to discard cells and large debris. The supernatant was then centrifuged at 10,000g for 20 min at 4°C to pellet microvesicles and other debris. To isolate EVs, the supernatant from previous centrifugation was ultracentrifuged 100,000g for 90 min at 4°C and the pellet (100K pellet) was washed in 12 ml of PBS and centrifuged again at 100,000g. Pellets were resuspended in 100-150 μL of PBS. All ultracentrifugation steps will be performed at 4°C using a SW41 Ti rotor (Beckman Coulter, Brea, CA, USA).

### Virus like particles (VLPs) preparation and collection

To produce Lentivirus-based virus like particle (VLPs), 293 T Lenti-X cells were transiently transfected on 10 cm Petri dishes with 8µg of pGagGFP plasmid, encoding the codon-optimized Gag protein fused to the green fluorescent protein (GFP) (30), using JetPrime transfection kit (Polyplus Transfection, Illkirch, France). After 48 h, cell culture supernatants were recovered, cleared from cellular debris by low-speed centrifugation, passed through a 0.45 μM pore size filter (Millipore Corporation, Billerica, MA, USA) and concentrated by ultracentrifugation for 2.5 h at 65,000 × g using a 20% sucrose cushion. VLPs were dissolved in 1× phosphate buffered saline (PBS, Gibco) and stored at −80°C until use.

### Flow Cytometric Acquisition of Beads, EVs and VLPs

All acquisition experiments were performed using a CytoFLEX LX flow cytometer (Beckman Coulter Life Sciences), equipped with five lasers (355, 405, 488, 561 and 640 nm wavelengths). This instrument allows to set the side scatter on Violet laser (V-SSC) in addition to the standard blue laser side scatter (B-SSC). Although PBS seems an obvious choice for a sheath fluid, PBS may increase the formation of calcium phosphate crystals in the fluidics system, which may cause additional background noise, clogging, or loss of laminar flow. Therefore, we used purified water as a sheath fluid. Anyway, scatter and all fluorescence parameters were set to log scale. We use two different fluorescent reference beads to fine-tune the instrument: Megamix-Plus FSC with the following diameters: 0.1 µm, 0.3 µm, 0.5 µm and 0.9 µm and Megamix-Plus SSC with following diameters: 0.16 µm, 0.20 µm, 0.24 µm and 0.5 µm (BioCytek Marseille, France). The beads were mixed (Megamix-Plus SSC/FSC) and diluted with PBS, following the instructions in the technical datasheet. Beads and all samples (EVs and VLPs) were diluted in PBS. Before starting the acquisition, the instrument underwent a thorough washing protocol. This procedure is crucial for minimising contaminants and reducing non-specific signals. Acquisition data were analysed by CytExpert v2.3 software.

### Flow Cytometry sorting of EVs

Sorting experiments were performed on MoFlo Astrios-EQ flow cytometer (Beckman Coulter), equipped with four lasers (405, 488, 561 and 640 nm wavelengths) and with a high-sensitivity forward scatter detector (488ex-FSC). Furthermore, the Astrios-EQ system is provided with inline sheath canister filters of 40 nm pore size. Forward scatter (FSC) detectors (FSC1 and FSC2) mounted the mask M1 and P1, respectively. Instrument were aligned with Rainbow QC Beads (Spherotech), while the detector settings for nanovesicle detection and fine-tuning adjustments of the alignment were made with Megamix-Plus SSC/FSC beads (BioCytek). The instrument acquisition software was the Summit v6.3.1. Analyses were performed with both Summit and FlowJo™ v10.9.0 Softwares.

### Statistical analyses

The results are shown as the means ± S.D. The data were evaluated for statistical significance using Student’s t-test. Any p value <0.05 was considered statistically significant.

## RESULTS

### Threshold settings for detection of fluorescent EVs

We previously described a promising approach for the study of fluorescent EVs (24, 31), where the evaluation of fluorescent EVs using a conventional flow cytometer (Gallios, Beckman Coulter) was described. The main strategy was to use their fluorescent emission rather than light-scattering detection, setting the threshold on fluorescence channel. This approach allows the detection of particles regardless of their size.

The evolution of technology led us to transfer this approach to more advanced instruments with enhanced performance, such as the CytoFLEX LX (Beckman Coulter). The initial step involved defining the sensitivity limit of the instrument. Since the CytoFLEX LX is equipped with the violet laser (405nm wavelength), we compared the resolution efficiency in separating small fluorescent beads gathered on violet side scatter (V-SSC) with the more common blue side scatter (B-SSC). Given that FSC is less sensitive to resolving EVs, a mix of two types of green fluorescent beads, Megamix Plus FSC and SSC, with dimensions between 100 and 900 nm, was employed as a reference to determine the size sensitivity limit of the cytometer. These reference beads enabled the establishment of both the threshold and gain levels, achieving a good compromise between sensitivity limits and the exclusion of background noise. The visualization of all the fluorescent beads was obtained by modulating the threshold and gain values on the FITC fluorescent channel (band pass 525/40), maintaining the background noise of the PBS barely detectable (<10 events/second), as shown in Figure 1A and B, both for V-SSC and the B-SSC. As shown in Figure 1C-F, the threshold set to fluorescence allows all beads to be detected on both V- and B-SSC channels, although the separation of the smallest beads (between 100 nm and 200 nm) is much more efficient on V-SSC compared to B-SSC (Figure 1C, D, red boxes). Furthermore, dot plots 1C and 1D confirm that small beads cannot be detected by their size on the Forward Scatter (FSC) parameter while V-SSC allows a better separation of compared to B-SSC (Figure 1E, F). Then, to identify fluorescent EVs, both threshold and gain were defined on the appropriate channel using non-fluorescent EVs (blank EVs) (Figure 1 G, H). Fluorescent EVs were purified from the cell culture supernatant of the A375M melanoma cell line pulsed with the green fluorescent fatty acid, BODIPY™ FL C16 (hereinafter as C16, ex/em 505nm/512nm), as previously described. The threshold and gain values on the fluorescence channel (FITC channel, bandpass 525/40) were adjusted until the background noise and blank EVs were barely visualized (<20 events per second, Figure 1G, H). Note that there is an increase in background noise above 400-500 nm. As shown in Figure 1I-N, the fluorescence distribution and number of C16-EVs per microliter using both thresholds (V-SSC and B-SSC) are very similar (Figure 1M, N), but a better resolution of C16-EVs was confirmed, as shown on the V-SSC channel (see red boxes).

**Figure 1.**
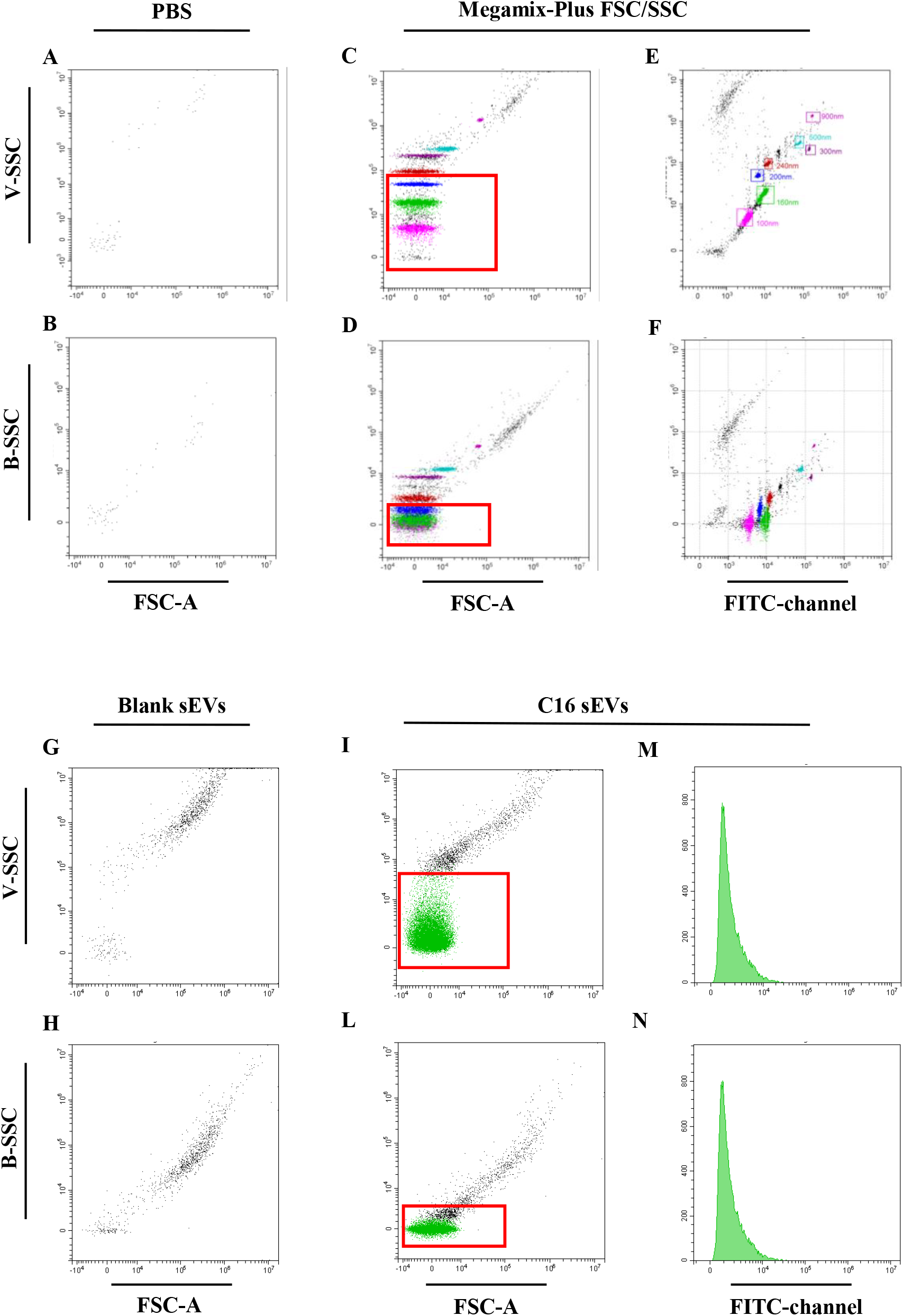
Comparison between Violet side scatter (V-SSC) and Blue side scatter (B-SSC) resolutions for fluorescent EVs detection. Panels A-B: FCM analysis of 0.22 μm filtered PBS acquired with threshold applied on FITC fluorescent channel (band pass 525/40). Panels C-E: Megamix Plus FSC/SSC polystyrene beads are used to define the resolution limit of the instrument and to compare the efficiency in separating smallest beads on violet side scatter (V-SSC) compared to the conventional blue side scatter (B-SSC). Threshold was applied on FITC fluorescent channel. Panels G-H: FCM analysis of unstained EVs released by A375M melanoma cell line (Blank EVs). Panels I-N: FCM analysis of fluorescent EVs released by A375M melanoma cell line pulsed with the green fluorescent fatty acid, BODIPY™ FL C16 (C16-EVs). Red boxes include beads between 100 nm and 200 nm in size.

An analogous acquisition strategy was applied to EVs secreted by cells pulsed with the orange-red fluorescent fatty acid, BODIPY™ FL C12 (referred to as C12-EVs, ex/em 558/568 nm). The threshold was applied to the PE fluorescent channel (band pass 585/42) in this case. To minimize background noise, we used PBS and blank EVs isolated from un-pulsed cells to set the threshold and gain values on PE channel. The histograms of Figure 2 display events recorded for PBS (Figure 2A and B) and blank EVs (Figure 2C and D) when applying the threshold on both FITC (on the left) or PE (on the right) channels. Figure 2E and 2F show the events recorded of C16- and C12-EVs, acquired with their respective threshold. As further control, we acquired C16- and C12-EVs by crossing the thresholds (Figure 2G and H).

**Figure 2.**
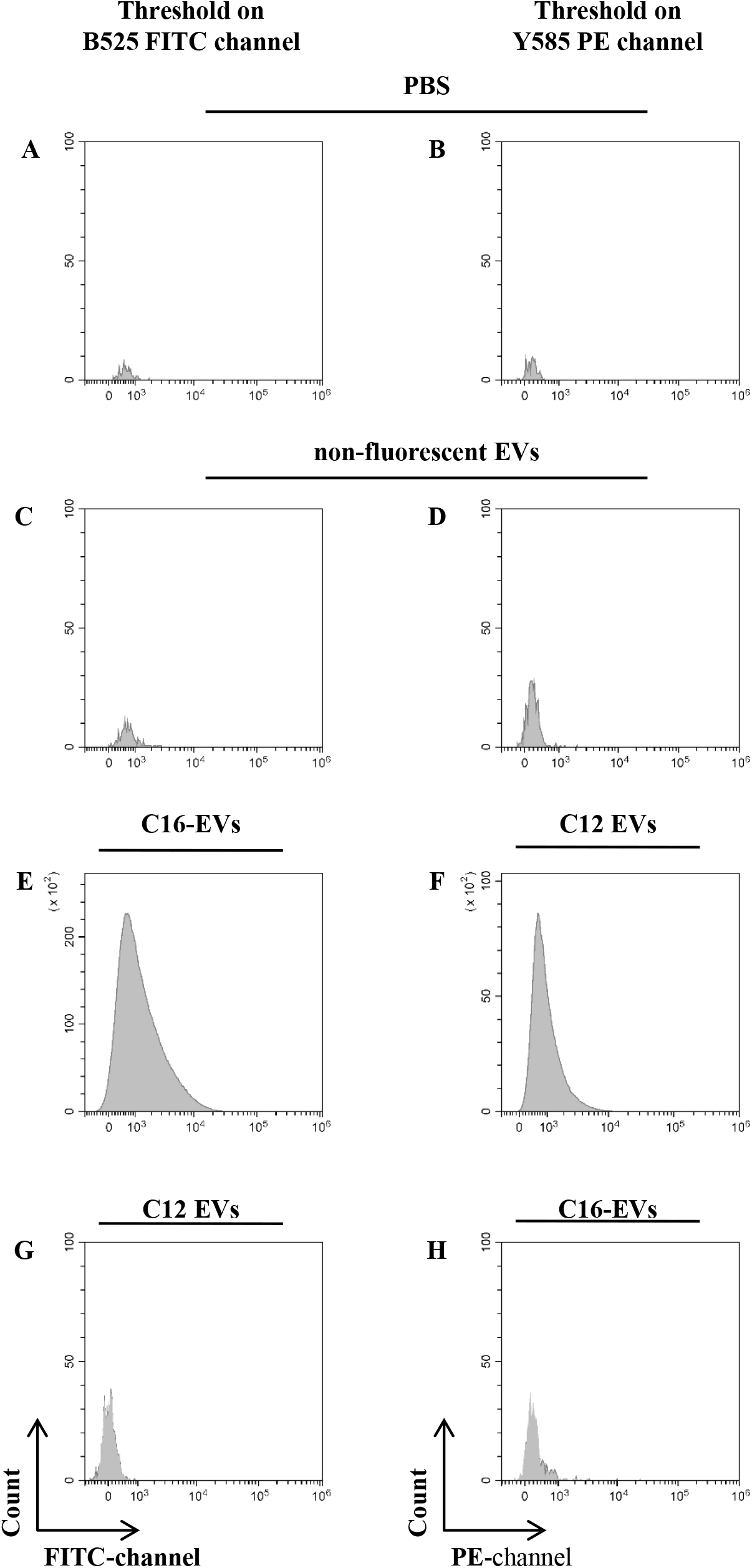
FCM profile of EVs purified from surnatant of A375 melanoma cell line. The histograms represent FCM analysis of 0.22 μm filtered PBS (A, B), unstained EVs, (C, D), C16-labeled EVs (E, H) and C12-labeled EVs (F, G). Fluorescent EVs were obtained pulsing cells with BODIPY™ FL C16 or BODIPY™ FL C12. Panel A, C, E, G: the threshold was applied on green fluorescent channel (B525 FITC channel). Panel B, D, F, H: the threshold was applied on orange fluorescent channel (Y585 PE channel). The threshold is set to reduce the background signal as much as possible (A, B, C, D) without losing the resolution of the appropriate fluorescent EVs (E, F). The histograms demonstrate the robustness of the analysis protocol: no events were found when C12-labeled EVs were analysed through green fluorescence threshold (G) as well as no C16-labeled EVs were found through PE fluorescence threshold (H).

These results demonstrate that thresholds applied to a specific wavelength can efficiently exclude other fluorescence emission associated with EVs during acquisition, and the presence or absence of signal in a specific acquisition protocol is not an artefact.

In conclusion, this approach for instrument setting allows for an easy and specific evaluation of fluorescent EVs in each sample, effectively minimizing background noise.

### Optimizing analysis conditions for high reliability FCM: Balance between Flow Rate, Abort Rate and Particles Concentrations

Quantification of EVs remains a challenging task and to address this, we have determined the optimal conditions for EVs-FCM experiments. High concentrations of EVs can result in a high number of missed events (high abort rate) and may lead to coincidence of two or more events, detected by the flow cytometer as a single event (swarm effect). To overcome this effect, it is very important to dilute the sample and/or reduce the sample flow rate, ensuring that only individual EVs exceed the threshold and contribute to the signals of the measured events.

In this context, the abort rate is a critical parameter to monitor in EVs-FCM experiments to obtain reliable results.

The ratio between flow speed and sample dilution must be carefully controlled. By modulating/lowering both flow rate and sample concentration, it is possible to find the best conditions to ensure linearity and high reliability on the number of acquired EVs.

Therefore, C16-EVs were serially diluted in 200 nm filtered PBS. We acquired C16-EVs at three different flow rates (slow 10 µl/min, medium 30 µl/min, fast 60 µl/min) and we applied the threshold on fluorescence to better detect and estimate the number of particles. The number of events per second, flow rate and the percentage of abort rate were recorded in each dilution and summarized in Figure 3. The results in graph 3A demonstrate a decrease in linearity when the abort rate exceeds 5% (dotted red line) and the events per second reach over 5,000.

**Figure 3.**
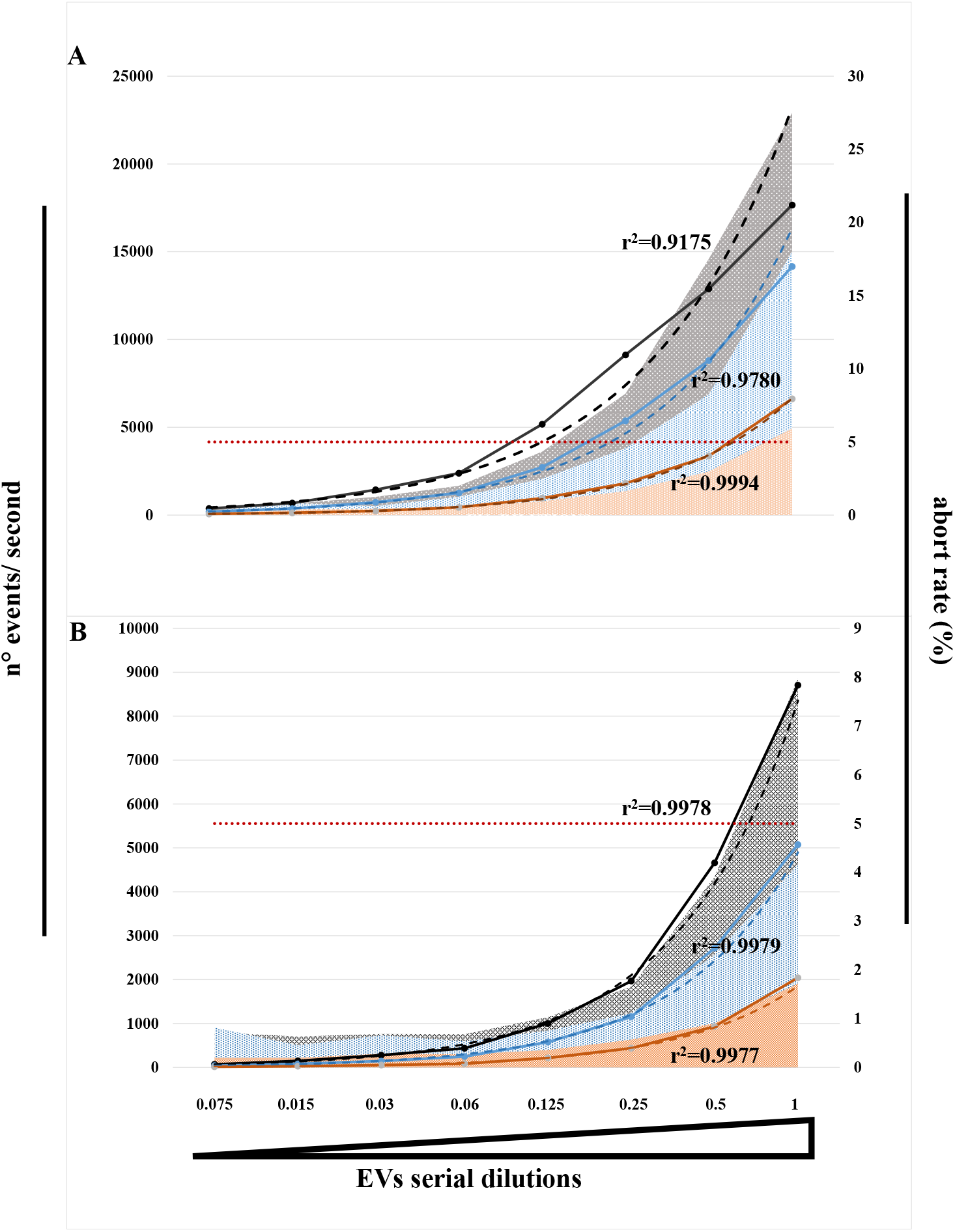
Effect of flow rate, abort rate and sample concentration on analysis accuracy. Several halving dilutions of C16-EVs was analysed at three different flow rates (red line-slow: 10µl/min, blue line-medium: 30 µl/min, black line-fast: 60 µl/min) and the number of events per second and the percentage of abort rate were evaluated. The dots on the continuous lines represent the number of events per second (y-axes on the left) at the indicated dilution (x-axes); dashed lines represent the fitted data with exponential model; shaded area represents the abort rate expressed as a percentage (y-axes on the right). A)High speed of acquisition (black line) combined with higher EVs concentration corresponds to a greater deviation from linearity (r^2^= 0.9175). Decreasing the flow rate increases the fitting of the model (r^2^= 0.9780) until slow flow rate (r^2^= 0.9994). Decrease in linearity is observed in all the flow rate conditions when the abort rate is exceeded by 5% (dotted line). B)The graph shows that it is possible to approach a value of r^2^ close to 1 even in conditions of high flow rate (black line), keeping the number of events/second below 5,000 and the abort rate below 5% (dotted black line). Results are representative of three independent experiments performed in duplicate.

High-speed acquisition (black lines) corresponds to a greater deviation from linearity (r2= 0.9175) where the maximum acquisition rate is around 18,000 events per second, while in medium- and slow-speed acquisitions the values of r2 are 0.9780 and 0.9994, respectively. However, the graph in Figure 3B shows that it is possible to approach a value of r2 close to one (r2= 0.9978) even under conditions of high flow rate (black line), keeping the number of events per second below 5,000 and the abort rate below 5%, regardless of the acquisition speed. The concomitant presence of high events per second, high speed and high abort rate represents the least efficient condition in terms of reliability.

### Flow cytometer set up and comparison between V-SSC and Fluorescence applied thresholds

As demonstrated previously (Figure1), the separation of beads of small size (between 100 nm and 200 nm) is much more efficient on V-SSC compared with B-SSC. Megamix-Plus SSC/FSC beads, a mix of beads that includes those below 200 nm, were used to draw a region including events below 200 nm (Figure 4A and B). The threshold applied on V-SSC allows the visualization of the background noise (Figure 4B) and the boundary between background signal and fluorescent EVs, necessary to draw the region, is shown in the dot plots of acquired of PBS (Figure 4C and D). C16-EVs were then analyzed applying thresholds on the fluorescent channel or V-SSC channel, showing similar number of events in both conditions (Figure 4E and F, red numbers in brackets).

**Figure 4.**
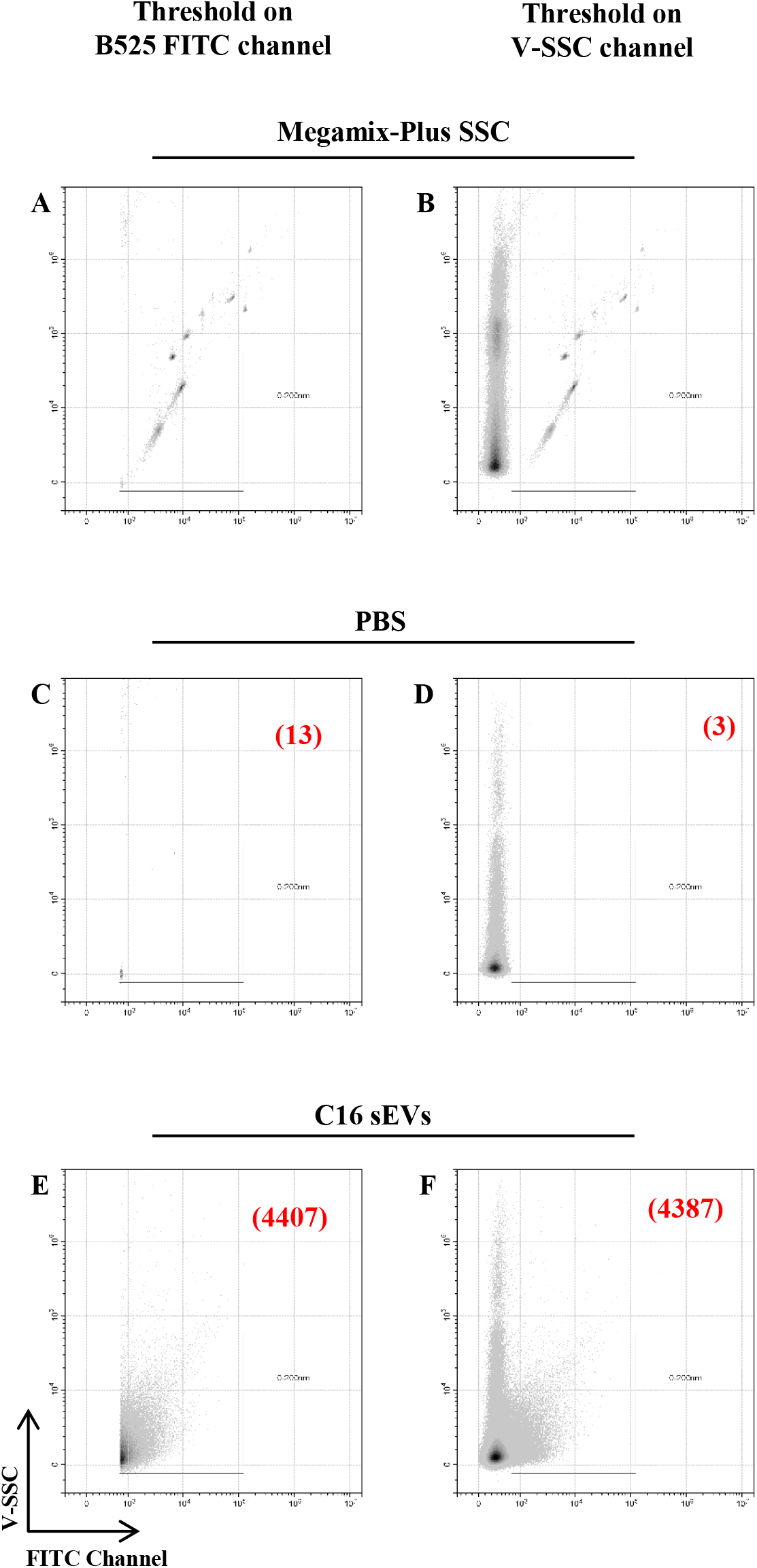
Comparison between V-SSC and Fluorescence applied thresholds. Representative density plots of reference beads (Megamix Plus FSC/SSC), analyzed applying the threshold on the fluorescence or on the V-SSC channels, are shown in A and B, respectively. PBS filtered at 0.22 µm was used to exclude the background noise (C, D). The black boxes were drawn to include the smallest sized beads (< 200 nm) and to exclude the background noise. The C16 EVs that fall in the black box were enumerated (E, F). The numbers in red, inside the dot plots C, D, E, and F, indicate the events per µl.

### Virus Like Particles acquisition

As shown previously, we performed a comparison between the two threshold strategies to establish overlapping results in terms of linearity and high reliability in the number of acquired EVs. A greater fluorescent intensity can allow better separation between background noise and fluorescent particles. Virus Like Particles (VLPs) used for these experiments were designed to carry the fluorescent fusion protein Gag-GFP. However, these VLPs lack the virulent components of their parent virus, eliminating the possibility that these virus particles could cause an active infection in an exposed individual. Their levels of fluorescence intensity are suitable for better separation of VLPs from background signals, allowing a reliable comparison of event counts by using the two different thresholds (Figures 5A-D). The fluorescent VLPs were acquired at same flow rate and the results are consistent with the previous observations (Figures 3 and 4). The number of VLPs are similar with both applied thresholds, although a full overlap is obtained with a flow rate ≤ 5,000 events per second (Figure 5E). A slight difference in the number of VLPs was observed above those values when setting the threshold on the fluorescent channel, although not significant (Figure 5E).

**Figure 5.**
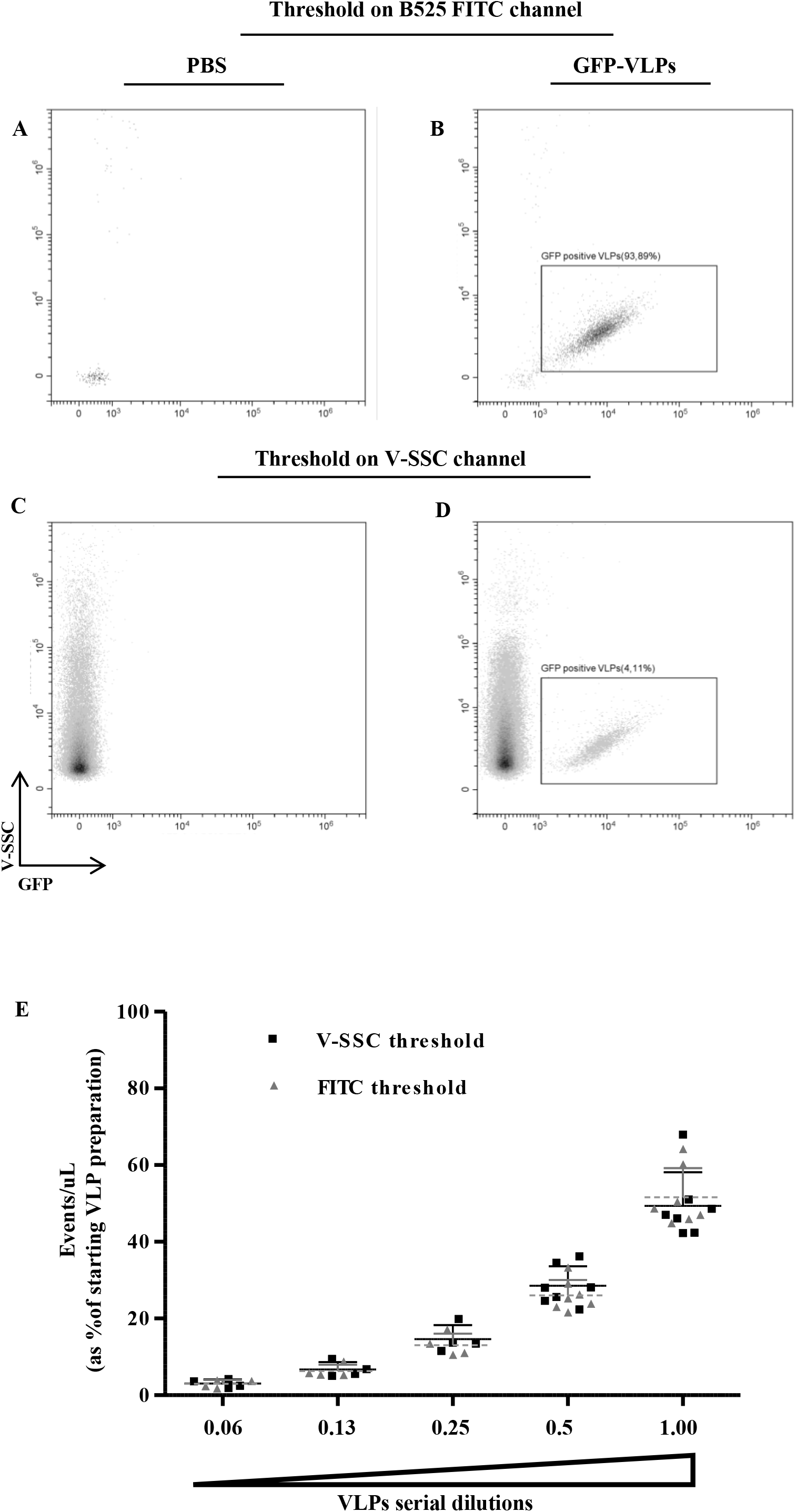
High reliability on the number of acquired nanoparticles. VLPs that carry fluorescent fusion protein Gag-GFP (GFP-VLPs) were quantified through the two different threshold strategies: A, B) threshold on green fluorescent channel; C, D) threshold on V-SSC channel. PBS buffer 0.22µm filtered (A, C) was used to determine instrument background noise. The black boxes were drawn to include fluorescent VLPs (< 200 nm) and to exclude the background noise. E) The graph summarizes serial dilution of VLPs count using the two different threshold strategies. The two series of data have a good fit (r^2^ > 0.995) as demonstration of the robustness of the method through counting with the two different threshold strategies. Count is expressed as percentage of starting VLPs preparation. Each experimental session was run with same acquisition settings (flow rate of 10 µl/min, threshold and gain value). The grey and black error bars indicate the standard deviation of samples acquired with the threshold applied on fluorescent or V-SSC channels, respectively. No significant difference was observed.

### EVs sorting

The most challenging goal is to demonstrate the feasibility of sorting EVs with high efficiency and purity, despite their small size. Since EVs show weak scattering properties and low antigen densities, having a reliable fluorescent tracer that allows clear identification of the vesicles to be separated from the background noise becomes an essential requirement. Recently, a method referred as “nanoFACS,” has been developed for the detection, analysis, and sorting of EVs and small viruses by flow cytometry (32). This method has been optimised for use with a customized MoFlo Astrios-EQ Cell Sorter (Beckman Coulter). We performed a similar experiment by using a MoFlo Astrios-EQ equipped with four lasers (405, 488, 561 and 640 nm), 70 μm nozzle, and with applied pressure of 60 psi. Although Morales-Kastresana et al have suggested a different approach (Morales-Kastresana et al., 2019), we have applied the threshold on 488-SSC (B-SSC) to allow a broader application of the procedure with commonly available instrumental configurations. As a basic strategy, we set a threshold value on B-SSC channel to visualize fluorescent reference beads smaller than 100 nm, as shown in Figure 6. Furthermore, to achieve optimal FCS separation, we tested several masks on dual forward scatter detectors, identifying the M1 and P1 masks as the best combination. Furthermore, we minimize vesicle aggregates and electronic background noise by a gating defined by plotting B-SCC Area channel versus B-SCC Width channel. In detail, as shown in Figure 6A, a region was drawn to exclude events with a higher B-SSC Width signal. The fluorescent beads Megamix-Plus FSC and SSC are shown in the dot plots (Figure 6 B, D) and histograms (Figure 6 C, E), respectively. Having defined the threshold value on the B-SSC channel and the corresponding voltage to reduce the PBS background noise to less than 300 events/sec, we confidently applied a simplified sorting procedure to sort EVs stained with different fluorescent tracers (Figure 7). In detail, two different productions of EVs, red fluorescent C12 and green fluorescent C16, were mixed and the sorting protocol was then applied. Of note, as shown in Figure 7B, EVs released by unstained cells were used as negative control (blank EVs). It is important to underline that, despite the refinement of the sorting protocol, it is not possible to eliminate the electronic background noise (quadrant Q4, Figure 7) (33). Therefore, the respective purities of sorted EVs must be evaluated considering exclusively the positive quadrants for the fluorescent signals (Q1, Q2, and Q3). Reanalysis of the C12-EVs and C16-EVs populations (Figure 7D, E) shown high levels of purity (98.4% Q1 and 96.6% Q3 respectively), while a non-significant number of EVs was identified outside specific gate (Q2+Q3 or Q2+Q1, respectively) evidencing the consistency of the method. Moreover, Figure 8 vividly illustrates the reliability of the method we have developed. In details, fluorescent EVs were sorted from cells incubated with both C16 and Ceramide, a red-fluorescent dye (λex 589nm / λem 616nm). Even though the two tracers, Ceramide and C16, were differently distributed on the EVs (Q1= 7.8% vs Q3= 83.1%, respectively, Figure 8A), the result of reanalysis shows a remarkable quality of the sorting (96.2% and 95.3%, respectively, Figures 8 B and C). Similar experiments were performed using cells transfected with a construct that carries a Nef^mut^-GFP fusion gene and incubated with C12 to obtain different EVs populations. The results, once again, demonstrate the feasibility of the protocol to sort EVs, allowing for C12-EVs and GFP-EVs purities of 94.7% and 97.3%, respectively (Figure 8 E and F). In conclusion, the high purity achieved after sorting, on specific EV populations, demonstrates the reliability of the developed protocol and the high quality of the procedure, regardless of the fluorescence channel and the characteristics of the fluorochrome.

**Figure 6.**
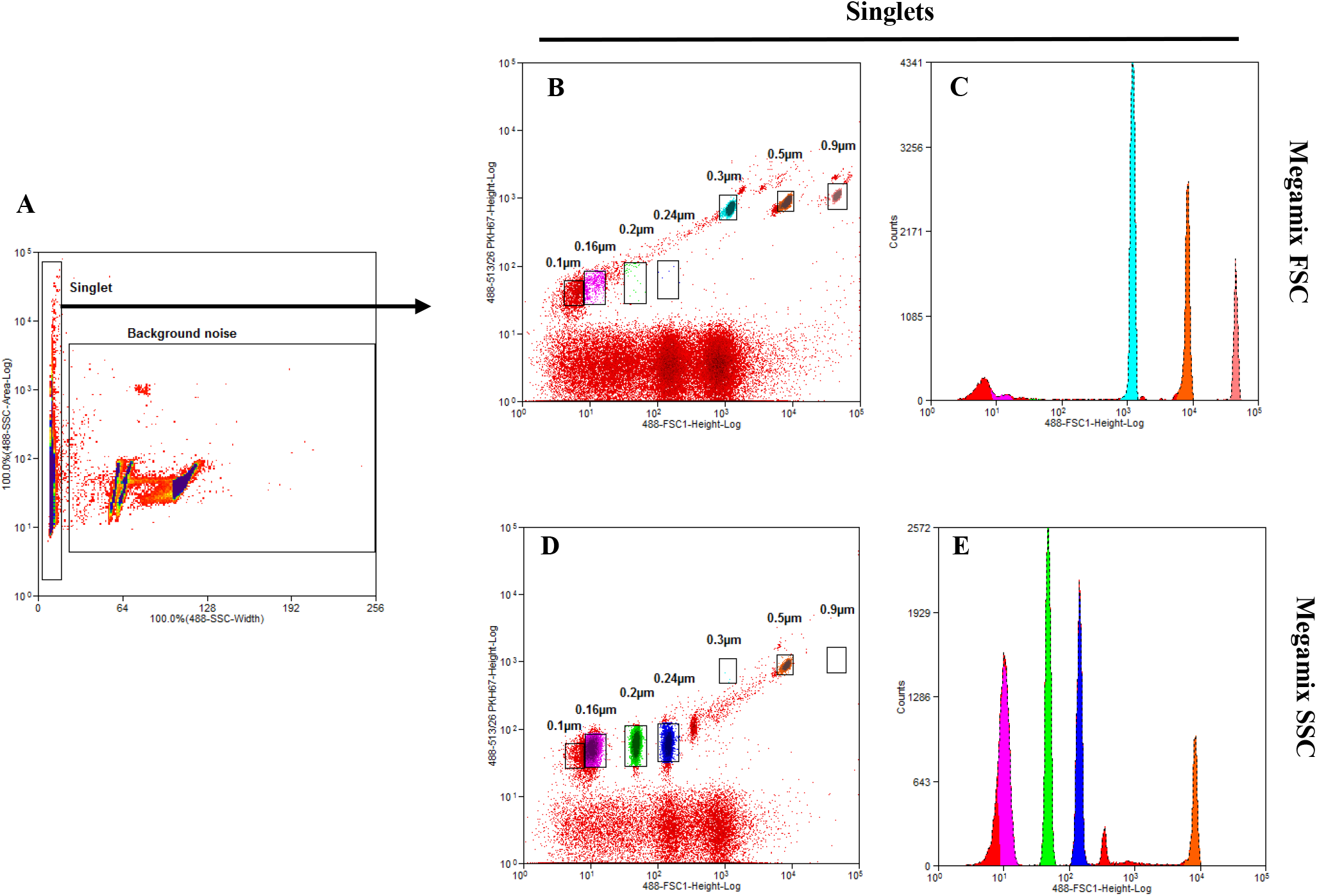
Gate strategy for EVs sorting. The B-SCC Area signal versus B-SCC Width signal allowed the exclusion of vesicle aggregates and background noise from the population of singlet particles (A). Representative dot plots and respective histograms of singlet gated Megamix-Plus FSC (B, C) and SSC (D, E) polystyrene beads with size between 100 nm and 900 mm (0.1µm, 0.16µm, 0.2µm, 0.24µm, 0.3µm, 0.5 µm and 0.9 µm). The analysis was performed on the events falling in the singlet region (see black arrow).

**Figure 7.**
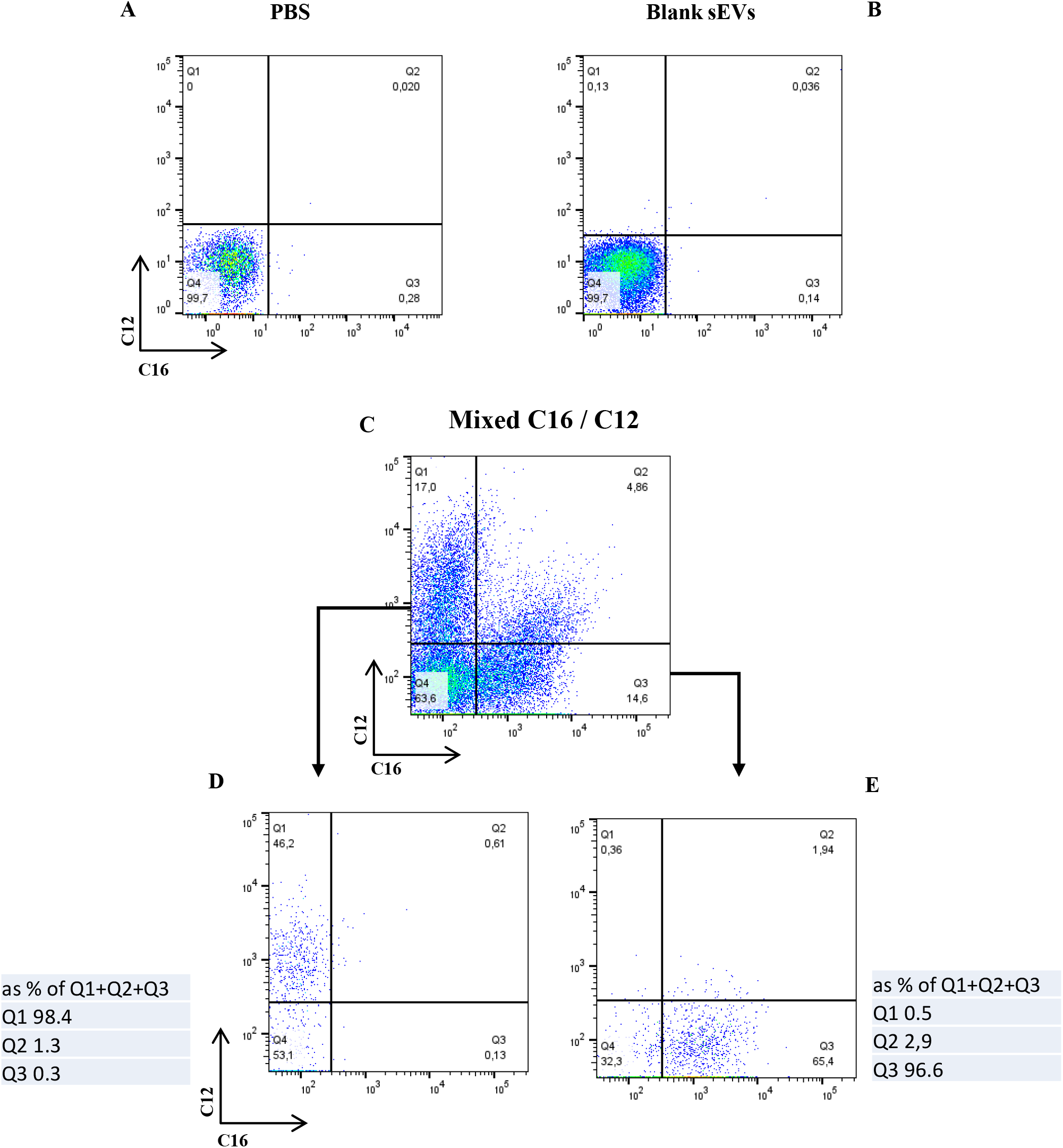
Sorting of fluorescent mixed C16 and C12 EVs. Representative dot plot of background noise assessed by PBS acquisition (A) EVs released by unpulsed cells were used as negative control for the fluorescent populations (B). Dot Plot of mixed C12- and C16-labeled EVs are shown in C. Reanalysis of sorted C12- and C16-labeled EVs are shown in D and C, respectively. The small tables on the left and on the right of D and F indicate the values of the positive quadrants for fluorescent signals (Q1, Q2, and Q3). The value of the positives is expressed as a percentage of the sum of the positive quadrants. The limitations of current flow cytometry technology cannot eliminate systemic background noise present in the quadrant Q4 that is excluded from the calculation of the positive EVs.

**Figure 8.**
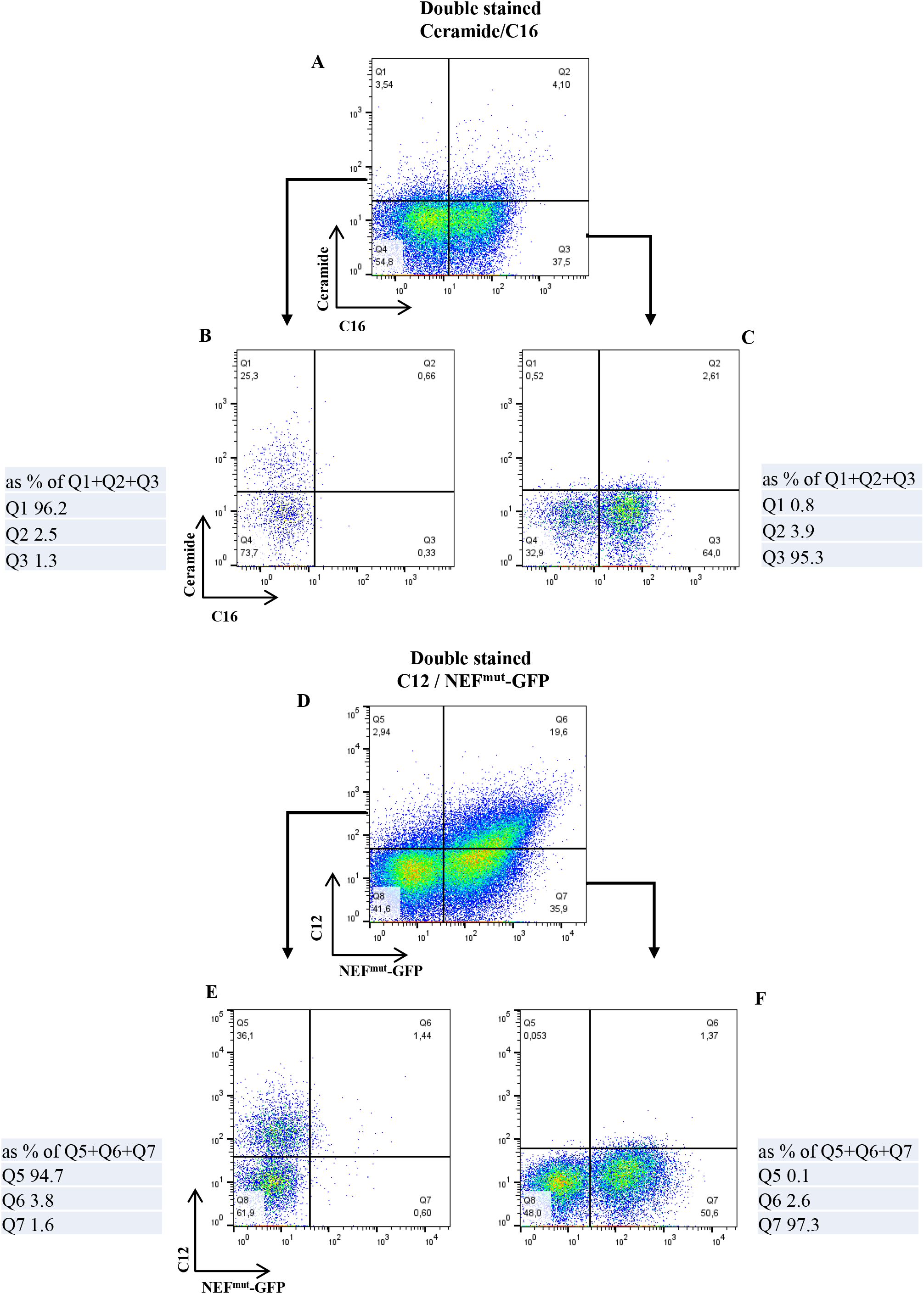
High resolution Sorting of Ceramide-C16 EVs and C12-Nef GFP EVs. The reliability of sorting tested on EVs sample obtained from cells incubated with C16 and Ceramide (A-C), and EVs sample released by cells transfected with a construct that carries a Nef^mut^-GFP fusion gene and incubated with C12 (D-E). The EVs population before sorting are shown on the dot plots A and D, respectively. Reanalysis of sorted EVs positive for a single fluorescence are shown on dot plots B, C, E, and F. The small tables on the left and on the right of the dot plots indicate the values of the positive quadrants for fluorescent signals (Q1, Q2, and Q3) and (Q5, Q6, and Q7). The limitations of current flow cytometry technology cannot eliminate systemic background noise in the quadrant Q4 and Q8 that are therefore excluded from the calculation of the positive EVs.

## DISCUSSION

EVs are a heterogeneous population of micro- or nano-membrane-bound particles involved in different biological functions. They play a role in communication between cells and organs, by carrying many classes of molecules that act as biochemical or molecular signals. They are characterized by a great heterogeneity in pathway generation, composition of bilayer, their cargo (4, 5). EVs can transport a variety of proteins, including growth factors, receptors, and cytokines, as well as lipids, nucleic acids, and metabolites, depending on their biogenesis. Furthermore, there is also a wide diversity in their size distribution: size range of EVs starts at ∼30 nm up to over 300 nm with a peak at a diameter of 100 nm (5, 34). EVs have garnered a lot of attention for their potential applications in diagnosis and their therapeutic potential (35). As well as EVs, VLPs are small particles of 20-200 nm that can be exploited as carriers for small molecules and are an attractive and versatile tool for the design of vaccines and therapeutics. Given the varying identities of these particles, a standardised approach is required to ensure reliable and accurate results for reproducible measurements and downstream applications. Minimal requests for reliable analysis technique are standardization of isolation methods; optimization of EV cargo analysis; accurate size-based quantification (36, 37). Various techniques try to respond to these methodological needs. For the purification of vesicles below 200 nm in size, size exclusion chromatography (SEC) and asymmetric flow field-flow fractionation (AF4) are commonly used isolation techniques. Quantification methods such as microscopy (scanning/transmission electron microscopy and atomic force microscopy), dynamic light scattering (DLS), nanoparticle tracking analysis (NTA), FCM, and small-angle X-ray scattering (SAXS) can be cited (10, 11, 38). FCM is a technique that offers several advantages, and as such, it has received significant attention. In detail, FCM is a low-time-consuming and high-throughput technique allowing high reproducibility, and in many cases, it may require low sample manipulation. The primary challenge is the detection limit, as it has mainly been used in cellular analyses or functional assays. Conventional FCM is not sensitive enough to detect EVs by morphological parameters. At nano-scale dimensions, there is not a linear relationship between light scatter and dimension, making the detection of EVs a challenge (39, 40). However, technological advances in new FCM have led to a significant improvement in the dimensional resolution, although there are still no tools capable of clearly identifying and quantifying EVs based solely on physical or morphological parameters. This study will optimize an FCM procedure for detecting nanoparticles with confidence. A straightforward protocol has been proposed for the reliable detection, quantification, and sorting of EVs released by cells in culture supernatants. The experience, gained from using the Gallios cytofluorimeter (31), has been successfully applied to the Cytoflex LX that boasts a wide linear dynamic range, resulting in improved resolution of dim and bright populations.

In conventional FCM protocols, a linear scale is used for light scatter measurements whereas a log scale is used for fluorescence signals, excluding some analyzes such as the cell cycle. In contrast, submicron particles are analyzed by non-conventional FCM settings, and scatter signals (FSC or SSC) are visualized with log scales. Furthermore, the threshold is set to remove signals primarily derived from debris and electronic background. Moreover, FCM instruments exhibit high background noise when calibrated for the analysis of small particles (approximately <200 nm). Using SSC downstream of other lasers, such as the violet laser (405nm) or yellow laser (561nm), will significantly improve instrument sensitivity, as suggested by Morales-Kastresana et al. (32). However, the chances of success increase when employing fluorescence signals, which are essential for identifying particles below 200 nm, separating them from background noise and debris (31, 33). Our study confidently presents results on the detection, quantification, and sorting of EVs. Identifying extracellular vesicles smaller than 200 nm (EVs) has been the first challenge. However, with the use of a fluorescent tracer, EVs can be accurately visualized even though conventional flow cytometers lack the capability to identify smaller particles based solely on their physical characteristics (Forward and Side Scatter). The second drawback was the selection of the parameter on which to apply the threshold. Our experience and the latest publications have highlighted the importance of fluorescence in visualizing nano-sized particles. Nevertheless, it is also feasible to utilize the enhanced sensitivity of SSCs when excited by lasers other than blue, thus eliminating the need for fluorescence. In our experimental setting, we used C16-EVs, a population of metabolically labeled EVs, approximately 80 nm in size (14) We decided to exclude direct-labeled EVs with fluorescent tracers or with specific fluorochrome-conjugated antibodies because standardized procedures to eliminate or reduce the presence of non-specific events are still under investigation (33, 41). As shown in Figure1, we demonstrated that we successfully detected fluorescent particles around 100 nm by using V-SSC. Additionally, we compared the number of events (C16-EVs) obtained by applying the threshold on the side scatter parameter with that applied on the fluorescence channel (Figure 4). In summary, our data confirm that applying the threshold on the violet laser (V-SSC) is more efficient in separating beads of different sizes than applying the threshold on the blue laser (B-SSC). Furthermore, we can clearly identify the EVs with a high level of correspondence by applying the threshold on the physical parameters, even without fluorescence. The main pitfall for a correct EVs quantification is the swarm effect and efforts should be made to avoid or greatly reduce multiple detections of EVs. Our studies have revealed that a harmonious balance of speed, concentration, and abort rate is essential in obtaining precise measurements of the particles being analyzed. In particular, the acquisition speed of individual events per time unit is significant in reducing the percent of abort rate (Figure 3). This parameter is crucial for establishing a window in which it is possible to define linearity conditions between flow rate, abort rate and EVs concentrations. In our working condition, the limit of abort rate to maintaining linearity has been set at 5%: by utilizing this value, we can enhance the acquisition speed while maintaining a linear regime (Figure 3B). Furthermore, to demonstrate the method’s robustness, we confidently enumerated C16-EVs using SSC or fluorescence thresholds, resulting in an excellent match of recorded events (Figure 4). The same protocol was successfully applied to analyze GFP-VLPs. Once again, the robustness of the method is highlighted, which exactly enumerates the particles analyzed with both the V-SSC and fluorescent thresholds (Figure 5). Therefore, the variability of the number of events counted is strongly influenced by the lack of control over the quantitative impact of background noise, which can be effectively reduced. Applying the threshold on the corresponding fluorescent channel for the quantitative evaluation of fluorescent EVs significantly reduces the background noise during acquisition compared to the background noise detected with standard protocols (SSC threshold). The use of a fluorescent threshold has already been demonstrated in the detection of EVs using conventional non-customized flow cytometer. New instruments with increased sensitivity enable better visualization of events below 200 nm. Although standardization of light scattering and fluorescence data between different flow cytometers remains a challenge, new instruments with increased sensitivity allow better visualization of events below 200 nm. Applying these suggestions has enabled the development of sorting strategies that utilize the threshold on the blue laser (B-SSC) with confidence. Our results demonstrate that sorting of EVs can provide reliable and accurate results even when using the B-SSC threshold. By separating vesicles labelled with different fluorescent tracers, a high level of purity can be achieved using cell sorters with well-defined parameters. As shown in Figure 7 and Figure 8 we reached a post sorting purity over 95%. Of note, it is fundamental to maintain the integrity of sorted particles and preserve their structure for downstream studies. In conclusion, our work emphasizes the importance of a pragmatic approach to instrument settings for the accurate identification and quantification of EVs.

## ACKNOWLEDGEMENTS

The authors acknowledge Dr. Francesco Manfredi (Istituto Superiore di Sanità, Rome) for providing the plasmid encoding Nef^mut^-GFP.

